# SpotLight proteomics identifies variable sequences of blood antibodies specific against deamidated human serum albumin

**DOI:** 10.1101/2023.04.09.536184

**Authors:** Jijing Wang, Susanna L. Lundström, Weiqi Lu, Yiqi Huang, Sergey Rodin, Roman A. Zubarev

## Abstract

Spontaneous deamidation of asparaginyl residues in proteins, if not repaired or cleared, can set in motion a cascade that leads to deteriorated health. Previously, we have discovered that deamidated human serum albumin (HSA) is elevated in blood of patients with Alzheimer’s disease and other neurodegenerative diseases, while the level of endogenous antibodies against deamidated HSA is significantly diminished, creating an imbalance between the risk factor and the defense against it. Endogenous antibodies against deamidated proteins are still unexplored. In the current study, we employed the SpotLight proteomics approach to identify novel amino acid sequences in antibodies specific to deamidated HSA. The results provide new insights into the clearance mechanism of deamidated proteins, a possible avenue for prevention of neurodegeneration.

## Introduction

Antibodies (Abs) are proteins produced by the humoral immune system to neutralize pathogens. They recognize the antigen via the complementarity-determining regions (CDRs) in the variable fragment antigen-binding (Fab) part of the antibody. To better understand how the acquired immunity works, we need to get a clear picture of how the amino acid sequences of Fab are selected in response to disease or infection.

An old paradigm in immunology posits that the Fab amino acid sequence selection is a completely random process, and thus the probability for two individuals both naive to a given immune challenge of having similar sequences raised in response to it will be vanishingly small [1, 2]. This old paradigm is being challenged by the new one postulating that, in a homogeneous group of individuals, the antibodies raised in response to a specific challenge to immune system, such as viral infection, will bear similarities in the sequences of the variable regions. This new paradigm has been supported by a number of recent studies [3–10]. Therefore, the fundamental task now pertinent to understanding the human immune system is to assemble a database of specific Fab sequences raised in response to different challenges. This work has already started, and information has been gathered on e.g. Alzheimer’s disease [11] and sarcoidosis [12] using the so-called SpotLight proteomics approach.

In the SpotLight approach, the antibodies are first purified from blood serum or plasma, and then digested by trypsin upon S-S bond reduction. The obtained tryptic peptides of the antibodies and co-purified proteins are separated by liquid chromatography and ionized by electrospray ionization. The ionized molecules are then analyzed by tandem mass spectrometry (MS/MS) employing two complementary fragmentation techniques, high collision dissociation (HCD) and electron transfer dissociation (ETD), which facilitates *de novo* amino acid sequencing [13, 14]. The obtained MS/MS spectra are first searched in the database of known sequences, and the unassigned peptides are *de novo* sequenced using appropriate software. Thus, obtained novel sequences that are common for most samples and strongly enriched in the patients compared to healthy controls are flagged as potential “hits”. These hits are further investigated in detail to remove possible false discoveries and compared with similarly obtained sequences from other diseases or infections [11, 12]. The validated sequences are then introduced into the database of antibody sequences.

It is of fundamental interest in immunology to understand which Fab sequences are expressed in response to which antigens. Knowing this will allow us to understand better how the human immune system works. One of the important antigens for which the human CDRs are yet unknown is isoaspartyl residue (isoaspartate, isoAsp). IsoAsp is a β-amino acid formed in proteins via deamidation of asparaginyl (asparagine, Asn) residue or, less frequently, isomerization of α-aspartyl (aspartate, Asp) residue. Deamidation is a facile, spontaneous, non-enzymatic and highly damaging posttranslational modification that turns every Asn residue to a potential time bomb inbuilt by nature into every protein [15]. Deamidation, if left unchecked, sets in motion a protein aggregation cascade that can ultimately lead to deteriorated health, ageing and neurodegeneration [15–18]. IsoAsp in blood proteins is therefore a risk factor for, e.g., AD [19] and other neurodegenerative diseases [20].

Previously, we discovered that deamidation may lead to the aggregation of human serum albumin (HSA), the most abundant protein in blood. Moreover, deamidation reduces the capacity of HSA to bind amyloid-beta (Aβ) peptide and phosphorylated tau (p-tau) protein, which affects Aβ and p-tau clearance enabled via HSA transporting them from brain to liver and kidneys [19]. The reduced clearance of Aβ and p-tau can ultimately cause neurodegeneration [21, 22].

One line of defense against deamidated HSA is its repair by PIMT enzyme in the liver; another one is the removal by endogeneous polyclonal antibodies (pAbs). However, the level of such pAbs in AD was found to be significantly diminished [15]. To test whether this is a common phenomenon in neurodegeneration, we have in a recent study purified IgGs from plasma samples of patients with AD, mild cognitive decline (MCI), frontotemporal dementia (FTD), vascular dementia (VaD), Parkinson’s disease (PD) and healthy controls (total n = 180) [20]. Using the artificially aged HSA (aHSA, isoAsp level ~ 60%) as an antigen, we have compared the levels of the IgGs against aHSA in these different disease groups with healthy donors. The results have revealed a significant reduction of anti-aHSA IgG levels in patients with AD, MCI and VaD (*P* < 0.01 in all cases), but possibly not in PD. These findings supported the notion that the deficit of anti-isoAsp antibodies in blood is a risk factor for neurodegeneration and necessitated studying such antibodies in greater detail.

One of the anti-isoAsp antibodies studied thoroughly is the murine monoclonal antibody of the IgG_3_ type that we have raised against a specific deamidated position in HSA [23]. We found that a single amino acid substitution in the heavy and light chain, respectively, can reduce the antibody specificity to isoAsp 1000-fold. Therefore, it is of considerable interest to find isoAsp-specific sequences in endogenous human antibodies.

Here, we embarked on such a mission with the following approach **(Figure 1).** Using artificially deamidated **HSA** (aHSA) as a bait, we purified specifically binding endogenous antibodies from healthy blood extracted from 100 individuals and uncovered the isoAsp-specific sequences with the SpotLight approach. As a control we used antibodies isolated in the same way but using fresh HSA (fHSA) as a bait. Then the unidentified MS/MS spectra were searched against a database containing both *de novo* sequences generated herein and Ig sequences deduced from patients in previous SpotLight studies [11, 12]. The obtained isoAsp-specific sequence information can help us better understand the clearance of damaged proteins from blood, and thus may show ways of preventing neurodegeneration.

**Figure 1.**
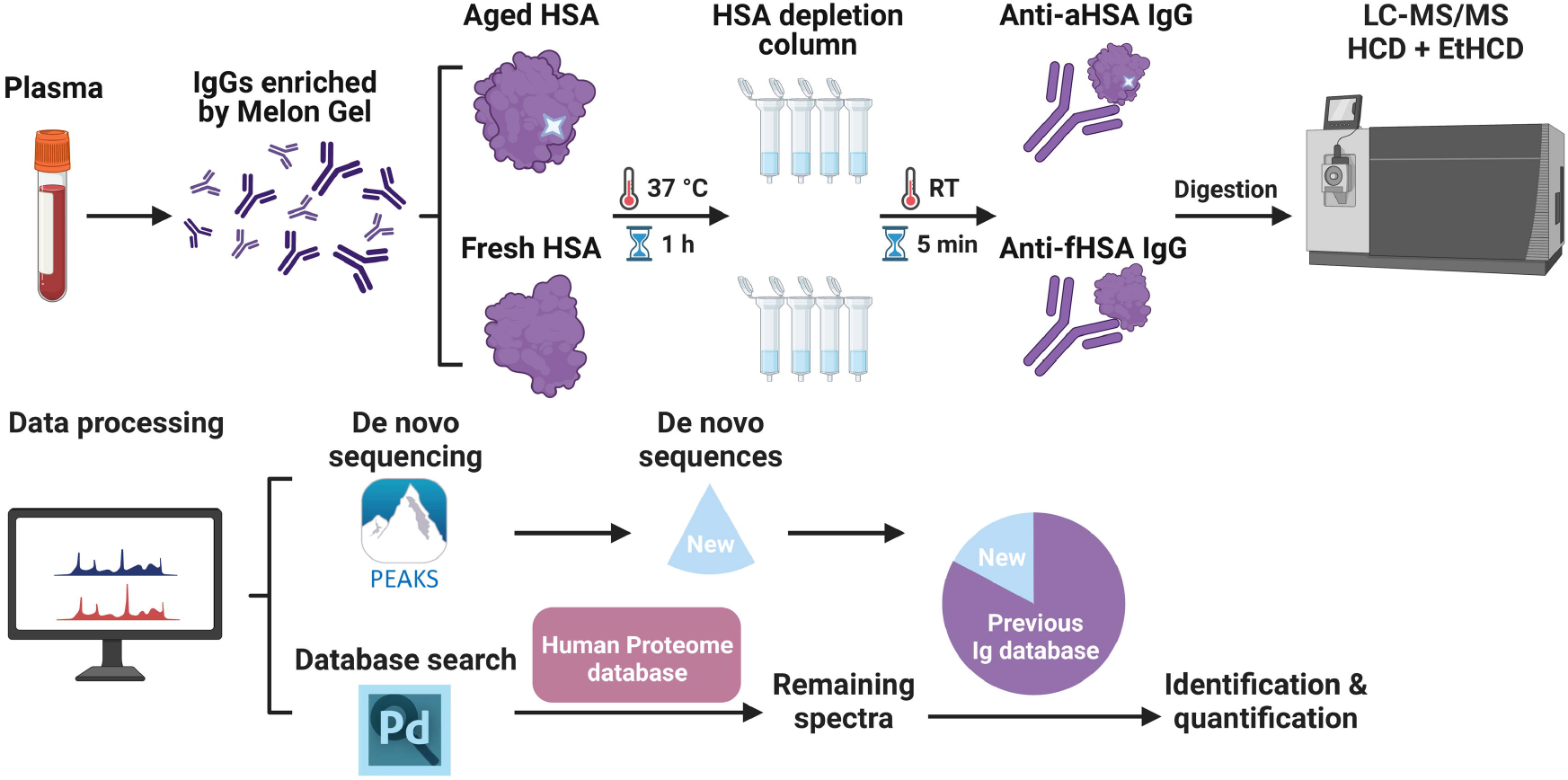
Schematic overview of the approach.

## Experimental Procedures

### Experimental Design and Statistical Rationale

Polyclonal antibodies were purified from blood using Melon Gel, incubated with aged HSA and fresh HSA (control), and the HSA-pAb complexes were isolated using an HSA depletion column. The complexes were then digested with a protease and analyzed by LC-MS/MS, with database search, *de novo* sequencing and abundance measurements. Statistically over-represented peptides for aged HSA novel sequences were selected.

### Patients and samples

Plasma from a cohort of 100 healthy blood donors (Age: 36 ± 11 years, Female: 41%) was obtained from ProMedDx Limited Liability Company. The samples were collected under a clinical study that had been reviewed by an Institutional/Independent Review Board (IRB) and/or independent Ethics Committee (IEC) according to the Declaration of Helsinki principles and the local regulations.

### Artificial aging of HSA

HSA (Sigma Aldrich) was incubated in 50 mM Tris buffer (pH 8.0) at 60 °C for 42 days, and then reduced, alkylated, digested, and analyzed by LC-MS/MS as described [2]. The isoAsp occupancy of all asparagine-containing peptides was determined; the peptide LVNTEFAK showed the highest occupancy (~ 60%).

### Purification of IgG from human plasma

The IgG antibodies were extracted from a pool of 100 healthy human serum samples using the Melon™ Gel IgG Spin Purification Kit (Thermo Fisher Scientific), with a tandem purification (one after another) performed in four replicates. Then the purified antibodies were incubated for 2 h at 37 °C with fHSA and aHSA, respectively, in a 1:1 molar ratio. The HSA-IgG complexes were immobilized on the albumin depletion column using the Pierce Albumin Serum Depletion Kit (Thermo Fisher Scientific) following the manufacturer’s protocol. The columns were washed gently three times by the binding buffer, and subsequently the HSA-antibody complexes were eluted from the column by 400 mM NaCl buffer.

### Digestion of purified HSA-IgG complexes and mass spectrometry analysis

The eluted HSA-IgG complexes were digested by Lys-C and trypsin (Promega) to peptides and analyzed by LC-MS/MS via HCD combined with electron-transfer/higher-energy collision dissociation (EThcD) fragmentation as in SpotLight analysis [11].

### Proteome Discoverer database search and quantitation

Peptide identification, assembly to proteins and label-free quantitation were performed using Proteome Discoverer 2.5 software (Thermo Fisher Scientific). First the MS/MS data were searched against the complete Uniprot human proteome reference database (UP000005640, 20509 entries) as well as a common contaminant database, with reversed sequences concatenated for FDR control. The 1% FDR was used as filter. The unidentified MS/MS spectra were then researched against a database containing Ig sequences (obtained from Uniprot) combined with *de novo* generated sequences obtained from both the samples herein and previous studies (433 entries, 516546 residues) [12]. The common contaminant database was also included in this second search. Peptide mass error tolerance was set at 10 ppm, while MS/MS fragment mass accuracy - at 0.02 Da in both searches. Cysteine carbamidomethylation was used as a fixed modification; methionine oxidation, asparagine and glutamine deamination were used as variable modifications for both identification and quantification. Trypsin was selected as enzyme specificity with maximum of two missed cleavages allowed. 1% FDR was applied as a filter for these data as well. Proteins identified as contaminants (except for serum albumin) as well as reversed protein sequences were removed. Proteins identified by at least 1 unique peptide and quantified with at least 2 peptides were kept. Quantification was performed by summing peptide abundances in individual mass spectra over peptide’s chromatographic peaks (i.e., by calculating the areas under the chromatographic curves). Proteins and peptides that were quantified in at least 4 out of 8 samples were kept for analysis. The abundances of Ig peptides and novel peptides were normalized to the total peptide abundance in all eight samples. The abundance of each protein was similarly normalized to the total protein abundance in all eight samples. When a peptide was observed in a given sample below the limit of quantitation range, instead of using zero as its abundance we used 1/10 of the lowest quantified value (determined for each protein or peptide specifically).

### PEAKs de novo sequencing

The raw MS/MS data were imported into PEAKS Studio 7.5 (Bioinformatics Solutions Inc), preprocessed via precursor mass correction, MS/MS de-isotoping and deconvolution, peptide feature detection, and then analyzed to generate a list of peptides. *De novo* sequencing was performed with 10 ppm mass error tolerance for precursors and 0.02 Da for fragment ions. Cysteine carbamidomethylation was set as a fixed modification, and methionine oxidation, asparagine and glutamine deamidation were marked as variable modifications. The *de novo* sequenced peptides were exported for the following statistical analysis.

### Fc-glycan profiling

Fc-glycan profiling was performed similarly to what has previously been described [24, 25]. Briefly, glycopeptides were identified by their characteristic retention times and accurate monoisotopic masses (within < 10 ppm from the theoretical values) of doubly and triply charged ions (IgG_1_: EEQYNSTYR, IgG_2_ or IgG_3_ [IgG_2/3_]: EEQFNSTFR and IgG_4_ or IgG_3_ [IgG_4/3_]: EEQFNSTYR (or EEQYNSTFR), respectively. Quantification of glycoforms was performed in a label-free manner using Quanti [26]. Glycopeptide ion abundances were integrated over respective chromatographic monoisotopic ion peaks (< 10 ppm from the theoretical values) at the charged states described above and within a ±1 min interval around the expected retention times. Glycoform abundances were normalized to total content (100%) of Fc-glycosylated IgG_1_ peptides, total content (100%) of Fc-glycosylated IgG_2/3_ peptides and total content (100%) of Fc-glycosylated IgG_4/3_ peptides, respectively.

### Statistical analysis

Univariate statistical analysis was performed using two-tailed Student’s t-test with equal or unequal variance depending upon the F-test, which resulted in *P*-values. FDR was calculated from the *P*-value and the number of tested variables (n = 36 for proteins and n = 537 for the Ig- and de novo-sequenced peptides) using the Storey Tibshirani method of multiple-hypothesis correction. FDR corrected *P*-values < 0.05 were considered significant.

## Results

### Peptide analysis

In total 536 peptides originating from immunoglobulins (Ig) or from *de novo* sequenced peptides with unknown origin were found in the anti-fHSA-IgG and anti-aHSA-IgG pools. After multiple-hypothesis correction, *P*-values were converted to false discovery rate (FDR), and FDR < 0.05 was taken as a threshold for statistical significance.

Of all peptides, 91 (17%) were significantly over-represented in terms of their abundances when aHSA was used as a bait compared to just 30 peptides (6%) with fHSA **(Table 1 and Figure 2).** Given that fHSA as a normal blood component should not trigger an immune response, while deamidated aHSA is immunogenic, this trend was expected. Notably, among the aHSA-specific peptides were not only Ig-Fab sequences (n = 43) and novel peptides (n = 21), but also sequences obtained from the conserved immunoglobulin regions (n = 27) **(Table 1).** Importantly, some of the enriched peptides formed groups exhibiting sequence similarity **(Table 2). A** full description of all peptides is given in **Supplementary Table 1.**

**Figure 2.**
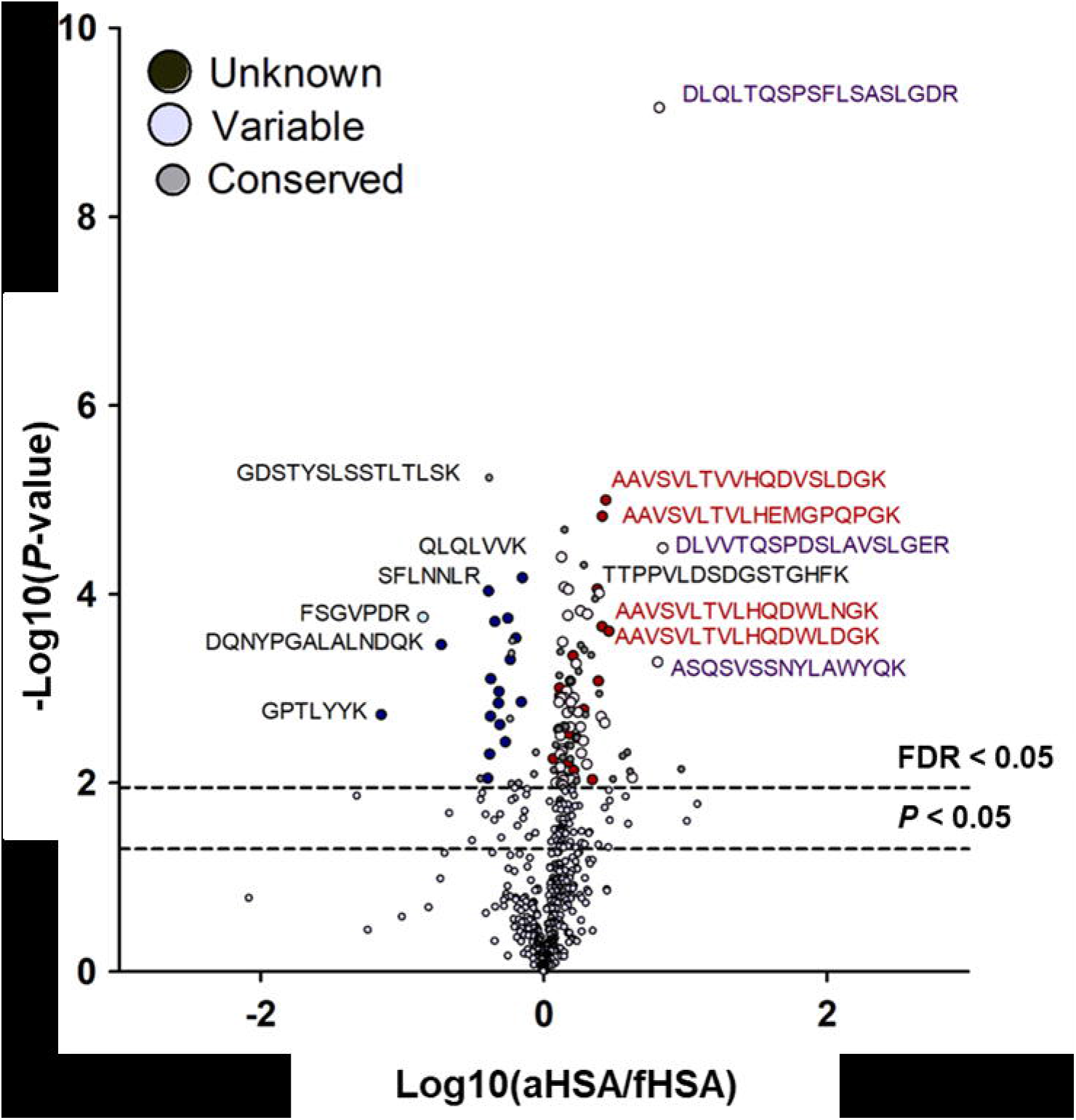
Volcano plot of the Ig-matching (Variable and Conserved) and *de novo* sequenced peptides (Unknown) with either aHSA specificity (positive x-values) or fHSA specificity (negative x-values).

### Comparison with SpotLight-derived specific sequences for neurodegenerative disorders (DLB and AD) as well as sarcoidosis

A total of 117 significant peptides from variable regions or unknown origin in this study were compared to those in two other Spotlight studies: 322 peptides from Study 1 (S1) related to Dementia with Lewy Bodies (DLB) and with Alzheimer Disease (AD) [11], and 426 peptides from Study 2 (S2) [12] on patients with two types of sarcoidosis and healthy controls. Of the total of 865 of such peptides in all three studies, 12 peptides were found in more than one study **(Table 3).** Notably, all these peptides were found enriched in aHSA complexes. Of these peptides, the most interesting are those originating from the CDR regions, such as the HV sequence LSCAASGFTFDDYAMHWVR and the KV sequence LLLYSASTLQSGVPSR from CDR1 and a CDR2, respectively. Previously the same peptides were found to be significantly elevated in bronchoalveolar lavage of sarcoidosis patients compared to healthy individuals. Also, three of the *de novo* sequenced peptides with unknown origin, YWGQGTLVTVSSASTK, WGQGTLVTVSSASTK and LLLSWASTR were both significantly elevated in the sarcoidosis patients. The new sequences WGQGTLVTVSSASTK and LLLSWASTR were also found to be elevated in the AD patients compared to DLB. The annotated MS/MS spectra of sequences listed in **Table 2** and **Table 3** are shown in **Supplementary Document.**

### Protein abundances

The identified peptides were grouped into proteins when the same protein sequence was identified with at least one unique peptide and quantified with at least two (not necessarily unique) peptides. In total, 36 proteins were quantified this way **(Supplementary Table 2).** As expected, the samples contained mostly immunoglobulin heavy constant gamma chains 1 to 4 (IgG_1_-IgG_4_) as well as the light kappa and lambda chains. The samples also contained Immunoglobulin heavy constant alpha 1 and 2 chains (IgA_1_ and IgA_2_) and well as Immunoglobulin delta heavy chain (IgD). Furthermore, several complement proteins as well as complement activating proteins were observed. Notably, some proteins were found to be significantly enriched in either fHSA or aHSA samples. Most noteworthy, IgA_1_ and complement 3 (one of the key proteins in the complement cascade) had a higher abundance in the aHSA enriched pAb samples. A full description of all proteins found in the samples is given in **Supplementary Table 2.**

### Deamidated proteins

Previously, products of asparagine deamidation have been found enriched in AD blood [13]. Hence, we included deamidation as a variable modification in the database search. Since deamidation rarely goes to completion, the match was accepted when both the modified and unmodified peptide were found. The abundances of deamidated peptides were first normalized to those of their unmodified counterparts and thus obtained deamidation occupancies were compared between the two sample types. The deamidation level was strongly increased in the aHSA enriched samples, especially in peptides from serotransferrin (TF), Beta-2-glycoprotein 1 (APOH), haptoglobin (HP) and hemopexin **(HPX) (Figure 3).** In total, 22 deamidated peptides were significantly enriched in the aHSA samples, and only one peptide – in the fHSA samples.

**Figure 3.**
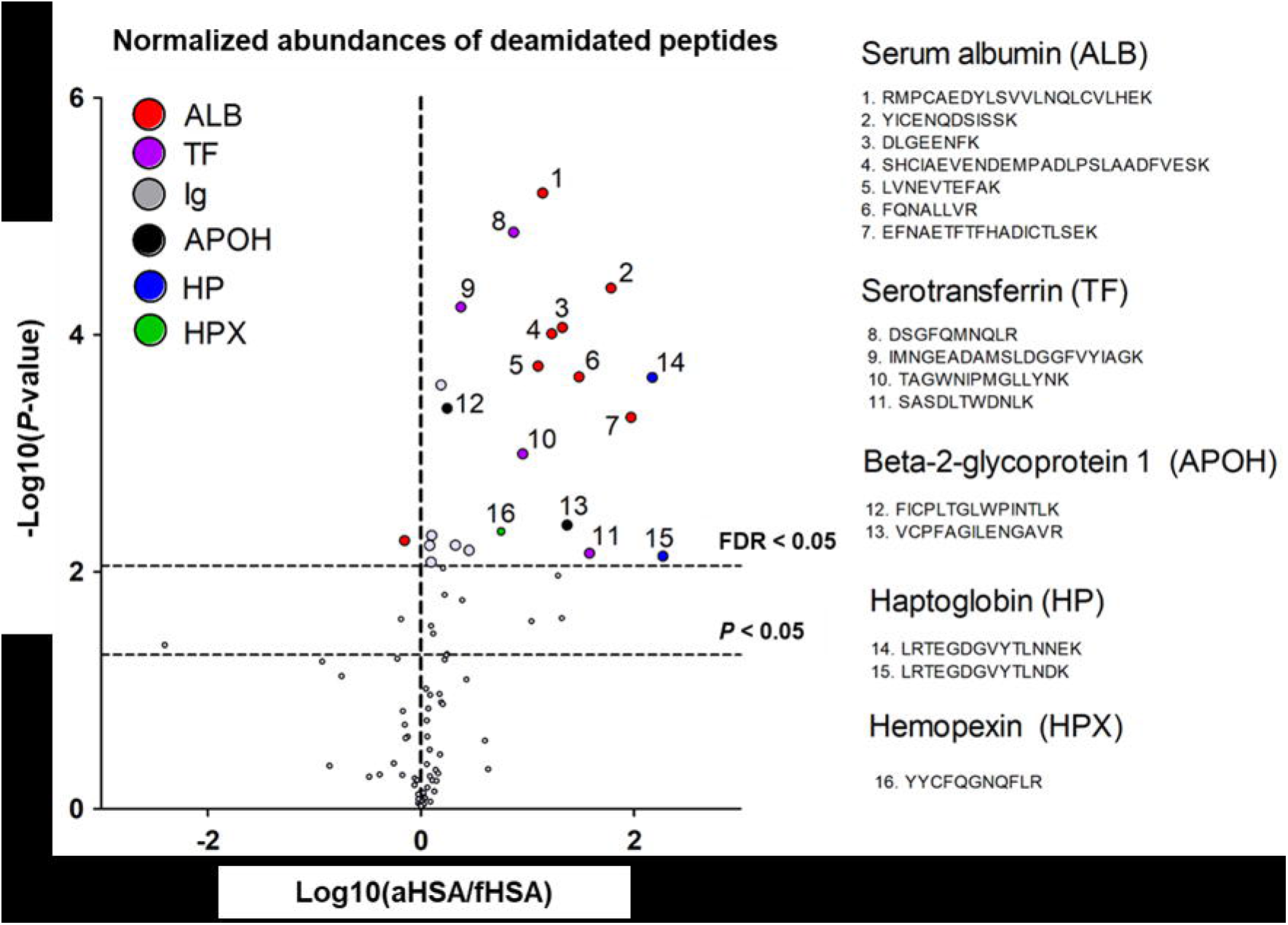
Volcano plot of the differences in the normalized abundances of deamidated peptides found in the fHSA- and aHSA-enriched pAbs.

Therefore, it appears that the pAb enriched against aHSA also exhibit specificity to other deamidated proteins. We investigated the position of these deamidation sites in the protein structures and found that a majority were located on the outer surface of the protein and thus are easily accessible for binding with antibodies **(Supplementary Figures 1-5). A** full description of all deamidated peptides found in the samples is given in **Supplementary Table 3.**

### Fc-glycosylotion patterns

Previously, we have shown that Fc-glycosylation patterns of pAbs of AD patients differ from those in healthy controls with a lower abundance of complex galactosylated and sialylated forms in AD [20]. Now we investigated whether the IgG Fc-glycosylation pattern would be different between pAbs enriched against fHSA and aHSA. A total of 18 glyco-peptides from IgG_1_ (EEQYNSTYR), 11 glycopeptides from IgG_2_ or IgG_3_ (IgG_2/3_, EEQFNSTFR) and 11 glycopeptides from IgG_4_ or IgG_3_ (IgG_4/3_, EEQ<u>F</u>NS<u>T</u>YR/ EEQYNSTFR) were identified, and their abundances determined. Overall, there were no distinct significant differences in the glycosylation pattern between the anti-fHSA-IgG and anti-aHSA-IgG **(Supplementary Table 4).** However, a difference was observed for sialylated forms of the IgG_4/3_ subtype which were discovered in significantly lower levels (*P =* **0.03)** in the anti-aHSA IgG.

## Discussion

In this study we set out to investigate the sequence differences between the pAbs specific to normal (fresh) human serum albumin compared to deamidated (aged) HSA. Previously, we have found that HSA deamidation leads to aggregation and loss of its carrying function. Importantly, deamidated HSA loses its ability to bind Aβ peptide and p-tau protein, which compromises the removal of these harmful molecules by their transfer from brain to liver and kidneys [19]. Thus, it has been postulated that an accumulation of deamidated HSA in blood is a risk factor for cognitive decline and neurodegenerative diseases [19]. Deamidated HSA and other deamidated blood proteins can be repaired in liver by PIMT enzyme and are also cleared from the system via deamidation-specific antibodies. It has been hypothesized that a bottleneck in this clearance process, resulting in accumulation of deamidated proteins in blood, can trigger neurodegeneration. This hypothesis has been strengthened in the recent studies [20, 27] in which the IgG levels of antibodies against aHSA were found to be significantly reduced in patients with AD, MCI and VaD compared to healthy donors.

Little is known regarding human antibodies against deamidation. Given that protein deamidation occurs constantly throughout life, it could be hypothesized that these antibodies are natural and present in a non-immunized organism starting from birth, similar to natural antibodies (nAbs) against Phosphorylcholine and Malondialdehyde [28]. Even though nAbs differ in function from adaptive antibodies, they tend to be polyreactive and bind to autoantigens and new antigenic determinants [29]. Noteworthy, the level of natural antibodies declines with age, which has been associated with age related illnesses such as cardiovascular disease and cancer [29, 30]. The most common nAb type is IgM, but both IgGs and IgAs can be of importance [31]. In this study IgA_1_ was distinctly enriched for aHSA, consistent with these pAbs to be native.

Furthermore, pAbs enriched against aHSA also reacted with other deamidated blood proteins, such as serotransferrin, beta-2-glycoprotein 1, haptoglobin and hemopexin. This observation may indicate that the anti-aHSA nAbs are cross-reactive with other deamidated epitopes and could possibly contain pan-isoAsp antibodies. The latter is of course a hypothesis that has a low probability, as pan-isoAsp antibodies are notoriously difficult to produce. For example, the murine 1A3 monoclonal antibody that we have generated against a specific deamidated site in aHSA did not show any cross-reactivity with other deamidated proteins [16].

Although the majority of the significant new or variable-region peptides were unique to this study compared to two other Spotlight proteomics studies **(Table 1, Figure 2),** some similar peptides were observed, all specific to aHSA **(Table 3).** In study S1 [11], more common peptides (four) were found elevated in AD than in DLB (two), consistent with HSA deamidation playing role in AD etiology. Among the peptides common with study S2 [12], seven were elevated in sarcoidosis patients compared to three peptides over-represented in healthy controls. This could be an indicator that deamidated HSA is a factor not only in neurodegeneration, but also in sarcoidosis. The cause of the latter disease is unknown but believed to be the body’s immune response to an unknown substance. More studies are needed to test the link between deamidation and sarcoidosis. Furthermore, the peptides with no sequence homology to either immunoglobulins or other proteins to which we refer in this study as “unknowns”, should be investigated further to determine if they are indeed CDR3 candidates and/or important for antigen binding.

Given that the fHSA is not immunogenic, the prevalence of aHSA-specific peptides (91 vs 30 specific for fHSA) was expected. It is possible that most of the 30 fHSA-enriched peptides are false positives, as the estimate is 27 false positives for 536 peptides detected at FDR = 0.05. If this is the case, the number of true positives in aHSA-enriched sample is close to 60. This result indicates, in line with other studies [25, 28], that the aHSA-enriched pool of pAbs exhibits significant degree of Fab sequence similarity, despite being extracted from serum of 100 individuals. Therefore, it is not unthinkable that this pool can be separated into several even more homogeneous groups of antibodies, perhaps dominated each by a single sequence, with subsequent *de novo* antibody sequencing.

## Conclusion

As *de novo* antibody technology is now relatively mature and accurate [16], it might be possible to derive in this way the sequence of an antibody against aHSA and, possibly, some other deamidated proteins. Such an antibody might have not only analytical potential, but also therapeutic prospects in preventing Alzheimer’s disease and other age-related disorders.

## Abbreviations

Abs: antibodies
AD: Alzheimer’s Disease
BAL: Bronchoalveolar lavage
C: Control
CDR: complementarity-determining region
DLB: Dementia with Lewy Bodies
ETD: electron transfer dissociation
EtHCD: electron-transfer/higher-energy collision dissociation
fHSA: fresh human serum albumin
FR: framework
HCD: high collision disso-ciation
HSA: human serum albumin
aHSA: aged human serum albumin
HV: heavy variable chain
IgG: immunoglobulin G
KV: kappa variable chain
LC-MS/MS: liquid chromatography coupled with tandem mass spectrometry
MG: Melon Gel enriched
NS: present but not significantly elevated
S: Sarcoidosis

## Data Availability Statement

All data generated or analyzed during this study are included in this publication and/or are available from the corresponding author on reasonable request. The raw MS proteomics data have been deposited to the ProteomeXchange Consortium (http://proteomecentral.proteomexchange.org/) via the PRIDE partner repository with the dataset identifier PXD039397.

## Supplementary Materials

This article contains supplemental data.

## Acknowledgments

The authors thank Marta Matlacz for the collection of part of the novel Ig sequences used in this study.

## Author Contributions

Conceptualization, R.A.Z.; methodology, R.A.Z., J.W., S.L.L., Y.H.; validation, J.W., S.L.L.; formal analysis, S.L.L., J.W., W.L. Y.H.; investigation, J.W.; resources, R.A.Z..; data curation, J.W., S.L.L., W.L.; writing—original draft preparation, J.W., S.L.L.; writing—review and editing, R.A.Z., S.R., Y.H.; visualization, S.L.L., J.W.; supervision, R.A.Z.; project administration, R.A.Z.; funding acquisition, R. A.Z., J.W. All authors have read and agreed to the published version of the manuscript.

## Table Legends

**Table 1.** Overview of the Ig-matching and *de novo* sequenced peptides of unknown origin found in the antibodies binding with fHSA and aHSA.

**Table 2.** Significantly over-represented peptides showing sequence similarity to other such peptides.

**Table 3.** Peptides from the variable region or *de novo* sequenced peptides of unknown origin that were significant in this study and at least one other Spotlight proteomics study (S1 and/or S2).

