## Supplementary material for "SpotLight proteomics identifies variable sequences of blood antibodies specific against deamidated human serum albumin": Tables

**Table 1.** Overview of the Ig-matching and de novo sequenced peptides of unknown origin found in the antibodies binding with fHSA and aHSA.

| **Type** | **Total found** | **Significantly enriched in** | |
| --- | --- | --- | --- |
|  |  | **fHSA** | **aHSA** |
| **Conserved** | 124 | 11 | 27 |
| **Heavy variable chain** | 101 | 0 | 20 |
| **Light variable chain** | 116 | 1 | 23 |
| **Unknown** | 195 | 18 | 21 |
| **Total** | 536 | 30 | 91 |

Abbreviation: Unknown, de novo sequenced peptides with unknown origin.

**Table 2.** Significantly over-represented peptides showing sequence similarity to other such peptides.

| **Enriched with** | **Sequence** | ***P*** | **Log10(aHSA/fHSA)** | **Origin** | **Region** |
| --- | --- | --- | --- | --- | --- |
| **fHSA** | AAFTSWTGGGAAASR | 0.02 | -0.38 | Unknown | - |
|  | VFTSWTGGGGR | 0.01 | -0.32 | Unknown | - |
|  | DQNYPGALALNDQK | 0.01 | -0.73 | Unknown | - |
|  | NYPGALALHHPNVAEK | 0.01 | -0.32 | Unknown | - |
|  | QLDLNVK | 0.005 | -0.26 | Unknown | - |
|  | QLQLVVK | 0.004 | -0.15 | Unknown | - |
| **aHSA** | AAVSVLTVLHEMGPQPGK | 0.003 | 0.41 | Unknown | - |
|  | AAVSVLTVVHQDVSLDGK | 0.003 | 0.44 | Unknown | - |
|  | AAVSVLTVLHQDWLNGK | 0.01 | 0.41 | Unknown | - |
|  | AAVSVLTVLHQDWLDGK | 0.01 | 0.46 | Unknown | - |
|  | FDPWGQGTLVTVSSASTK | 0.03 | 0.13 | Unknown | - |
|  | YWGQGTLVTVSSASTK | 0.005 | 0.26 | Unknown | - |
|  | VEDTAVYFCAR | 0.05 | 0.13 | HV | FR3 |
|  | SEDTAVYYCAR | 0.05 | 0.13 | HV | FR3 |
|  | ASDTAMYYCAR | 0.03 | 0.26 | HV | FR3 |
|  | AGDTAVYYCAR | 0.02 | 0.28 | HV | FR3 |
|  | LSCAASGFTFSSYGMHWVR | 0.04 | 0.12 | HV | FR1/CDR1 |
|  | LSCAASGFTFSSYWMHWVR | 0.02 | 0.26 | HV | FR1/CDR1 |
|  | LSCAASGFTFSSYWMSWVR | 0.04 | 0.30 | HV | FR1/CDR1 |
|  | LSCAASGFTFSSYAMSWVR | 0.02 | 0.18 | HV | FR1/CDR1 |
|  | LSCAASGFTFSSYSMNWVR | 0.01 | 0.28 | HV | FR1/CDR1 |
|  | LSCAASGFTFDDYAMHWVR | 0.005 | 0.31 | HV | FR1/CDR1 |
|  | LSCAASGFTFSDYYMSWIR | 0.02 | 0.40 | HV | FR1/CDR1 |
|  | LSCTASGFTFGDYAMSWVR | 0.02 | 0.16 | HV | FR1/CDR1 |
|  | LSCAASGFTFSNAWMSWVR | 0.004 | 0.39 | HV | FR1/CDR1 |
|  | ATGIPDRFSGSGSGTDFTLTISR | 0.01 | 0.14 | KV | FR3 |
|  | FSGSGSGTDFTLTLR | 0.004 | 0.14 | KV | FR3 |
|  | DLQLTQSPSFLSASVGDR | 0.05 | 0.08 | KV | FR1 |
|  | DLQLTQSPSFLSASLGDR | 0.003 | 0.81 | KV | FR1 |
|  | ALQMTQSPSSLSASVGDR | 0.03 | 0.13 | KV | FR1 |
|  | ELVLTQSPGSLSLSEGPR | 0.01 | 0.16 | KV | FR1 |
|  | ELVLTQSPGTLSLSPDGR | 0.01 | 0.11 | KV | FR1 |
|  | ELVLTQSPGTLSLSPGDR | 0.02 | 0.11 | KV | FR1 |
|  | VVLTQSPATLSLSGPER | 0.03 | 0.11 | KV | FR1 |
|  | LVLTQSGPTLSLSPGER | 0.02 | 0.11 | KV | FR1 |

Abbreviation: Unknown, de novo sequenced peptides with unknown origin; HV, heavy variable chain; KV, Kappa variable chain; Region, position in the variable sequence; FR, framework region, CDR, complementarity determining region.

**Table 3.** Peptides from the variable region or *de novo* sequenced peptides of unknown origin that were significant in this study and at least one other Spotlight proteomics study (S1 and/or S2).

| **Sequence** | **Type** | **Variable region** | **Significantly elevated in** | |
| --- | --- | --- | --- | --- |
|  |  |  | **AD or DLB (S1)** | **S or C** |
| LSCAASGFTFDDYAMHWVR | HV | FR1/CDR1/FR2 | - | S (BAL) |
| AGDTAVYYCAR | HV | FR3 | - | S (BAL) |
| TEDTAVYYCAR | HV | FR3 | NS | C (BAL) |
| AEDTAVYYCAR | HV | FR3 | AD (serum) | C (BAL) |
| LSCAASGFTFSSYGMHWVR | HV | FR1 | - | S (BAL) |
| VEDTAVYFCAR | HV | FR3 | NS | C (BAL) |
| ALQMTQSPSSLSASVGDR | KV | FR1 | DLB (MG) | - |
| DLVMTQSPDSLAVSLGER | KV | FR1 | DLB (MG), AD (serum) | NS |
| LLLYSASTLQSGVPSR | KV | FR2/CDR2/FR3 | - | S (BAL) |
| YWGQGTLVTVSSASTK | Unknown | - | - | S (BAL) |
| WGQGTLVTVSSASTK | Unknown | - | AD (serum) | S (BAL) |
| LLLSWASTR | Unknown | - | AD (MG) | S (BAL) |

Abbreviation: HV, heavy variable chain; KV, kappa variable chain; FR, framework; CDR, complementarity-determining region; NS, present but not significantly elevated; AD, Alzheimer’s Disease; DLB, Dementia with Lewy Bodies; MG, Melon Gel enriched; S, Sarcoidosis, C: Control, BAL, Bronchoalveolar lavage.
