## Supplementary material for "SpotLight proteomics identifies variable sequences of blood antibodies specific against deamidated human serum albumin": 1. Supplemental materials

**Supplementary Table 1:** List of Ig-peptides and unknown peptides found in the samples – *Excel file*

**Supplementary Table 2:** Proteins found in the samples – *Excel file*

**Supplementary Table 3:** List of the ratio between deamidated peptides/normal peptide in the samples *–* *Excel file*

**Supplementary Table 4:** Relative abundance of Fc-glycoforms on different IgG-Fc types *–* *Page 2*

**Supplementary Document:** Annotated MS/MS spectra of sequences listed in Table 2 – *PDF file*

**Supplementary Figure 1:** Volcano plot of the proteins found in the samples *–* *Page 3*

**Supplementary Figure 2:** Structure of human serum albumin (HSA) from Protein Data Bank (PDB, ID: 1e78) and positions of identified peptides *–* *Page 4*

**Supplementary Figure 3:** Structure of serotransferrin (TF) (PDB, ID: 6d04) and positions of identified peptides *–* *Page 5*

**Supplementary Figure 4:** Structure of Beta-2-glycoprotein 1 (APOH) (PDB, ID: 4jhs) and positions of identified peptides *–* *Page 6*

**Supplementary Figure 5:** Structure of hemopexin (HPX) (PDB, ID: 1qjs) and position of the identified peptide *–* *Page 7*

**Supplementary Figure 6:** Structure of haptoglobin-hemoglobin complex (PDB, ID: 4f4o) and position of the identified peptide *–* *Page 8*

**Supplementary Table 4.** Relative abundance of Fc-glycoforms on different IgG-Fc types.

|  | **Fc-glycoform** | **fHSA** | **aHSA** | ***P*** |
| --- | --- | --- | --- | --- |
|  | Gal (-) | 33 ± 12 | 34 ± 9 | 0.8 |
| **IgG_1_** | Gal (+) | 59 ± 9 | 58 ± 6 | 0.9 |
|  | Sia (+) | 8 ± 7 | 8 ± 3 | 0.9 |
|  | Gal (-) | 53 ± 3 | 53 ± 3 | 0.8 |
| **IgG_2/(3)_** | Gal (+) | 47 ± 3 | 47 ± 3 | 0.8 |
|  | Sia (+) | 0.1 ± 0.1 | 0.2 ± 0.2 | 0.2 |
|  | Gal (-) | 59 ± 5 | 52 ± 9 | 0.2 |
| **IgG_4/(3)_** | Gal (+) | 41 ± 5 | 48 ± 9 | 0.2 |
|  | Sia (+) | 3 ± 1 | 2 ± 1 | 0.03 |

Abbreviation: Gal (-), without galactose; Gal (+), with galactose; Sia (+), with sialic acid.

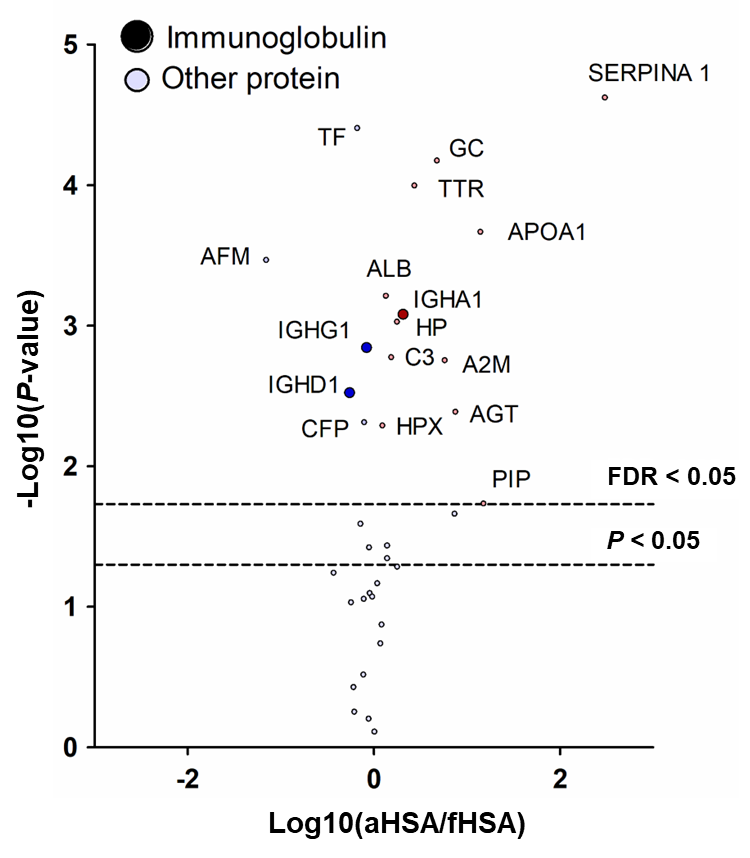

**Supplementary Figure 1:** Volcano plot of the proteins (denoted by their gene names) found in the samples. Proteins enriched in aHSA-specific samples show positive Log10(aHSA/fHSA) values.

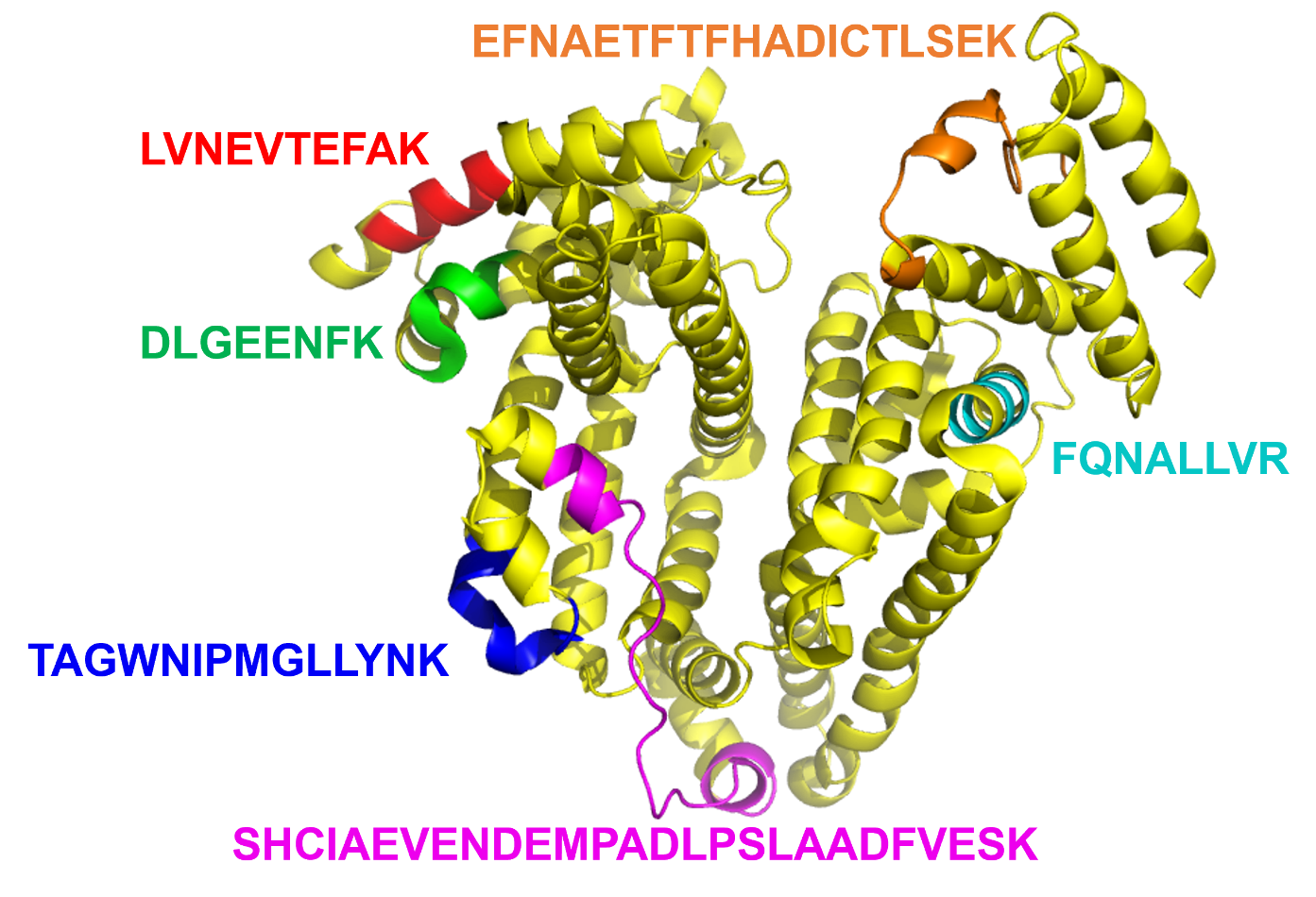

**Supplementary Figure 2:** Structure of human serum albumin (HSA) from Protein Data Bank (PDB, ID: 1e78) and positions of identified peptides.

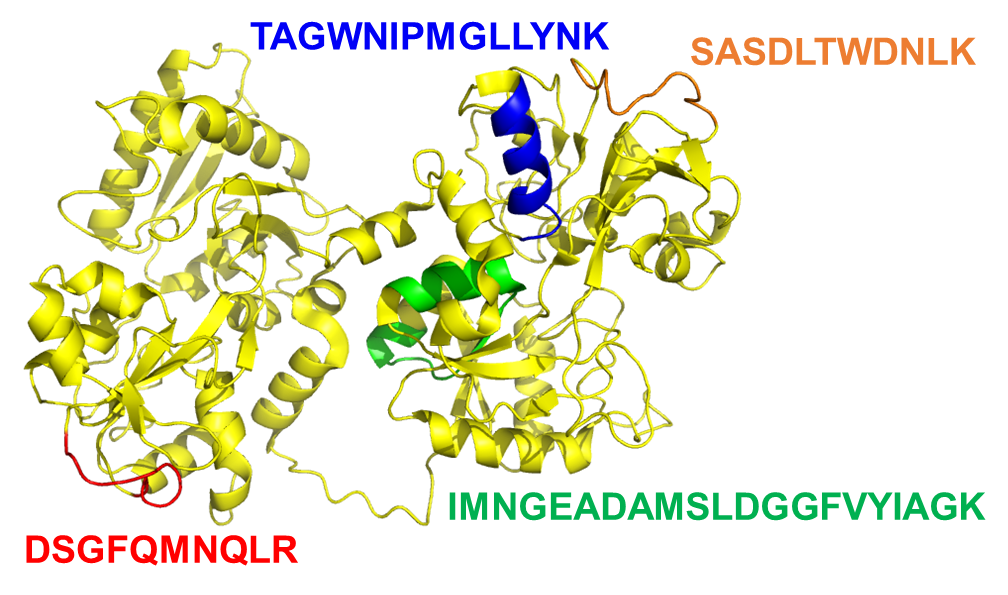

**Supplementary Figure 3:** Structure of serotransferrin (TF) (PDB, ID: 6d04) and positions of identified peptides.

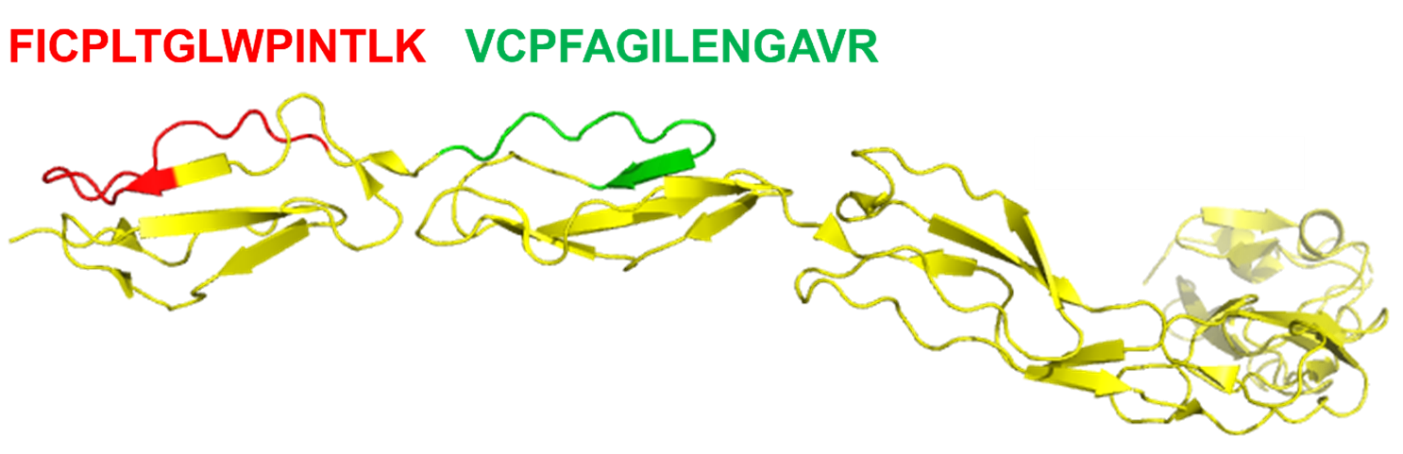

**Supplementary Figure 4:** Structure of Beta-2-glycoprotein 1 (APOH) (PDB, ID: 4jhs) and positions of identified peptides.

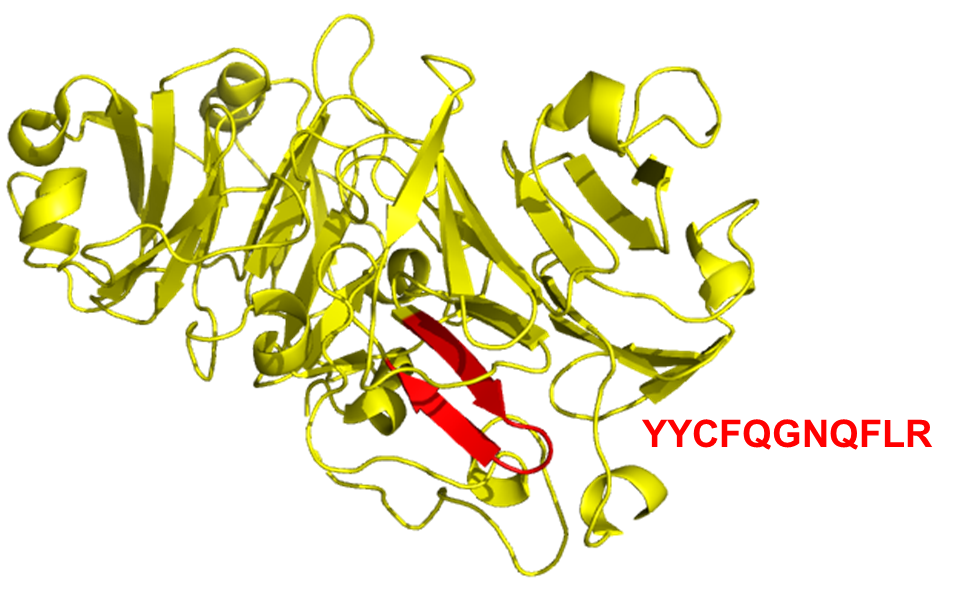

**Supplementary Figure 5:** Structure of hemopexin (HPX) (PDB, ID: 1qjs) and position of the identified peptide.

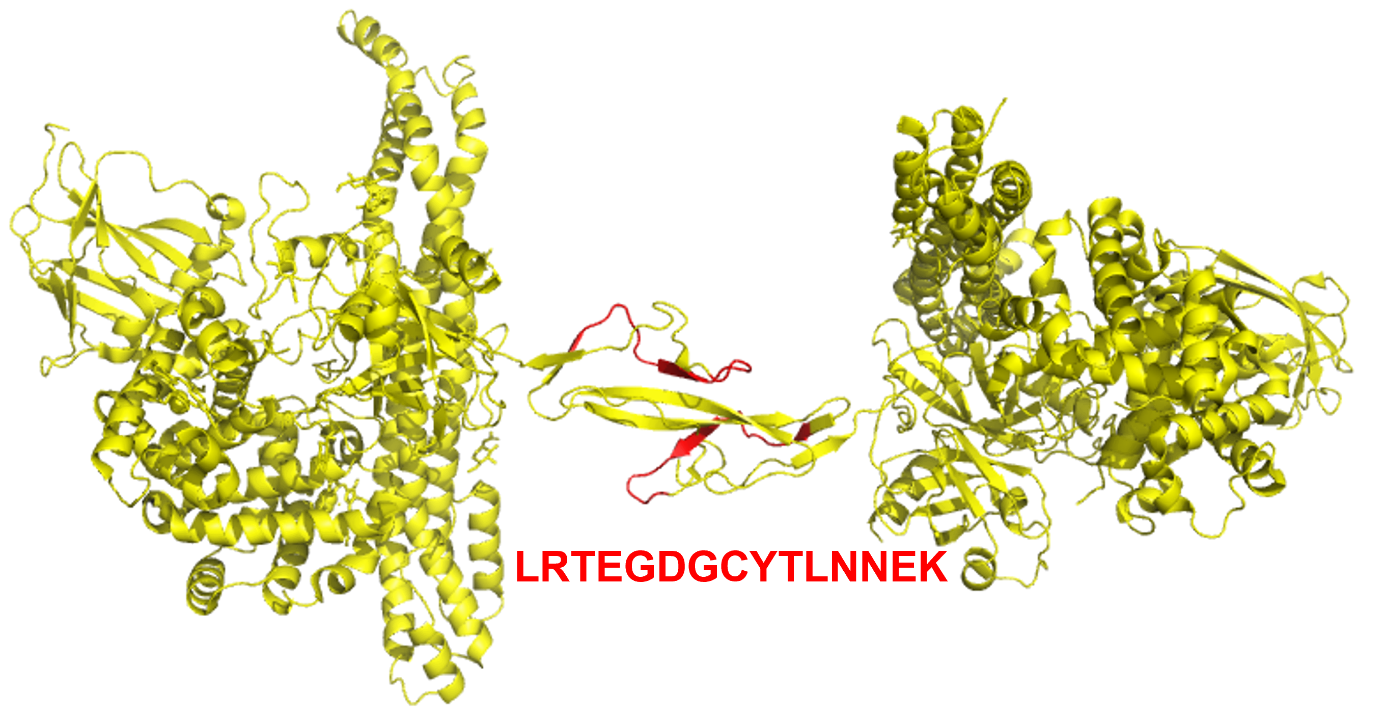

**Supplementary Figure 6:** Structure of haptoglobin-hemoglobin complex (PDB, ID: 4f4o) and position of the identified peptide.
