## Supplementary material for "SpotLight proteomics identifies variable sequences of blood antibodies specific against deamidated human serum albumin": 3. Supplementary Document

### Annotated MS/MS spectra of sequences listed in Table 2

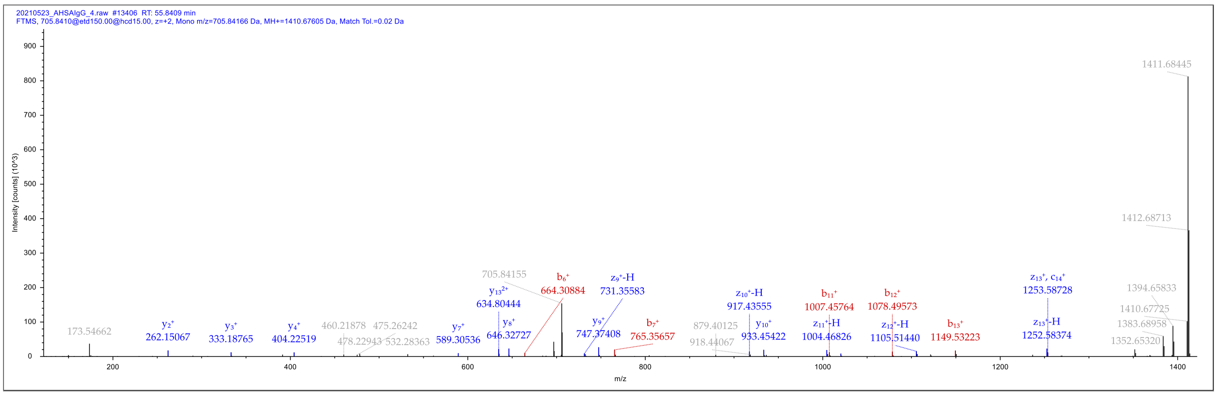


**AAFTSWTGGGAAASR**

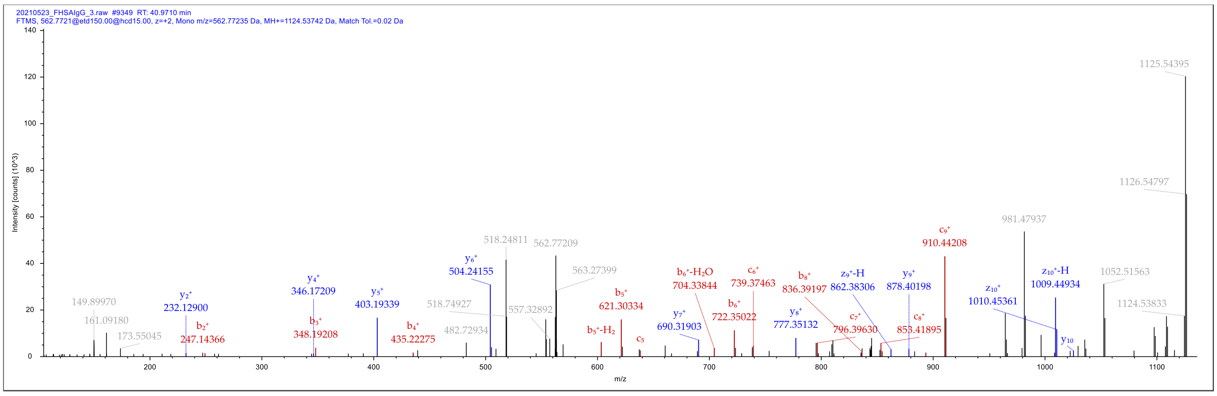


**VFTSWTGGGGR**

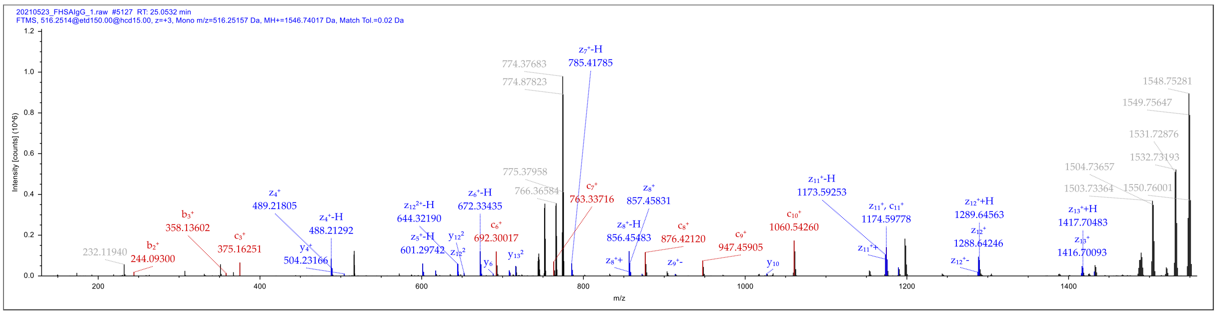


**DQNYPGALALNDQK**

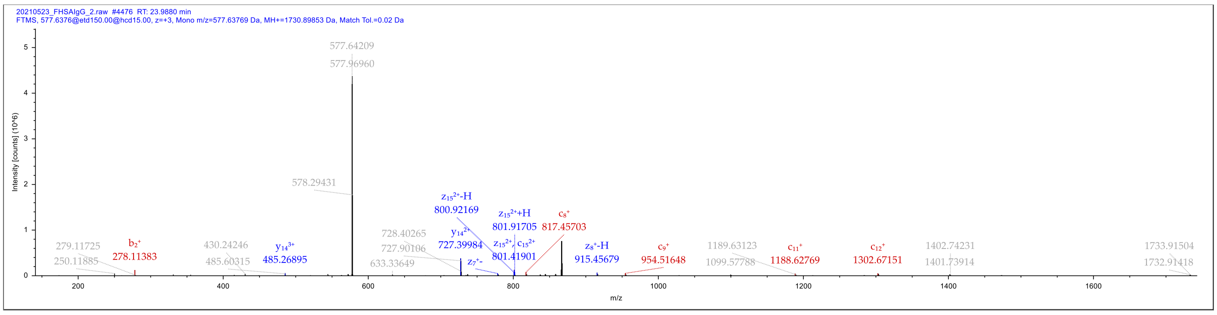


**NYPGALALHHPNVAEK**

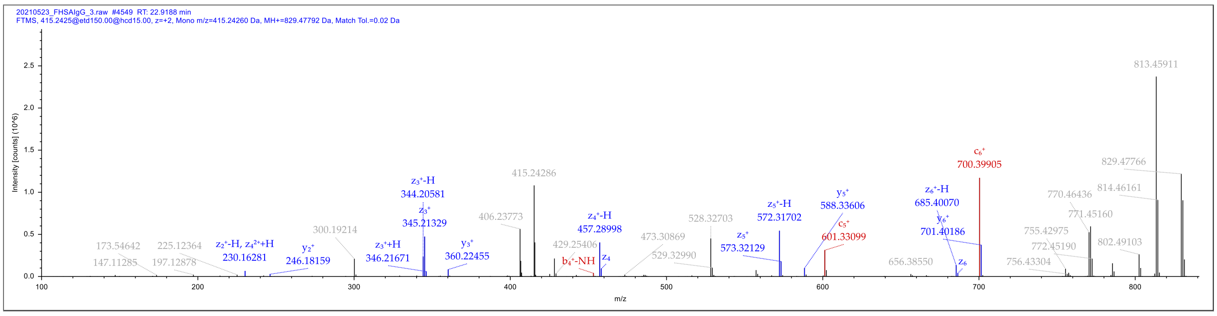


QLDLNVK

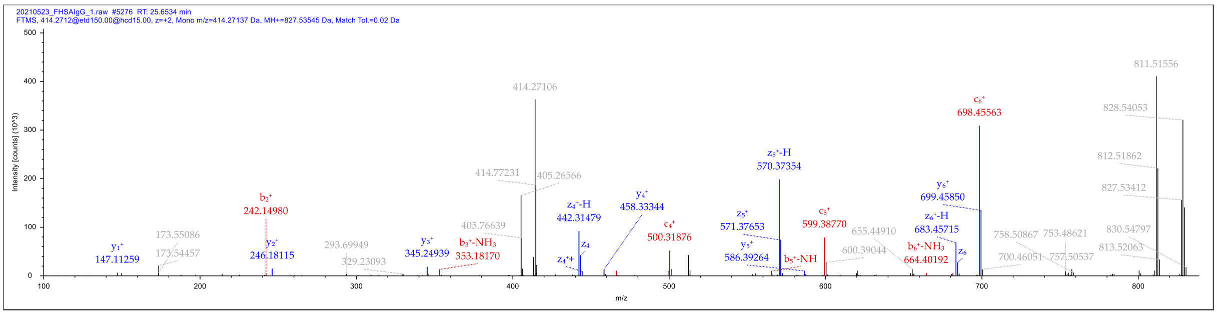


QLQLVVK

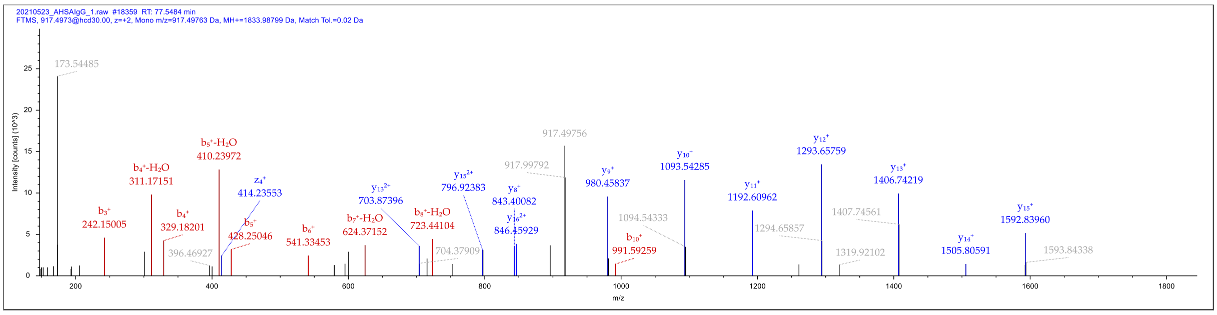


AAVSVLTVLHEMGPQPGK

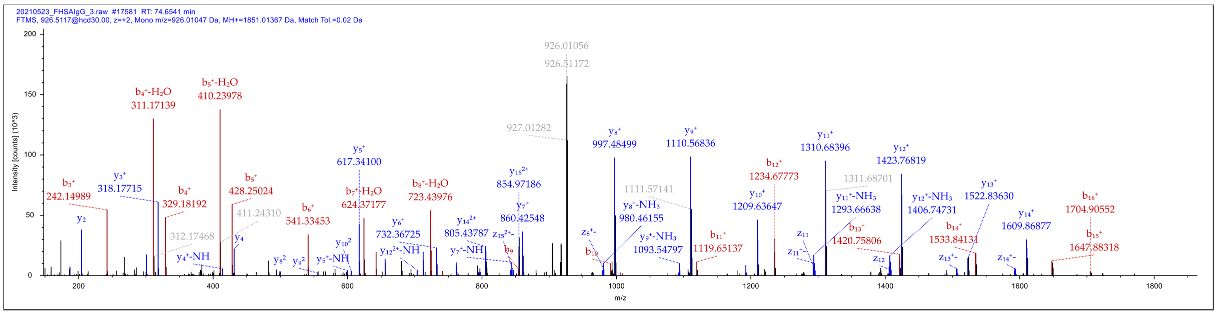


AAVSVLTVVHQDVSLDGK

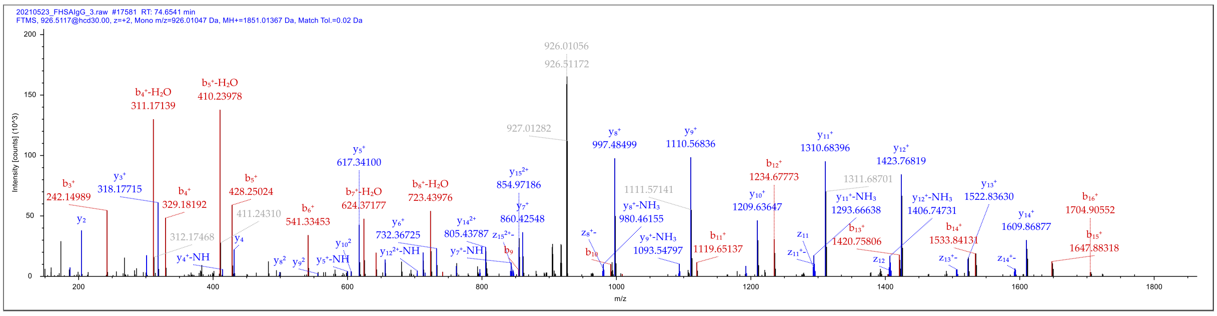

#### AAVSVLTVLHQDWLNGK

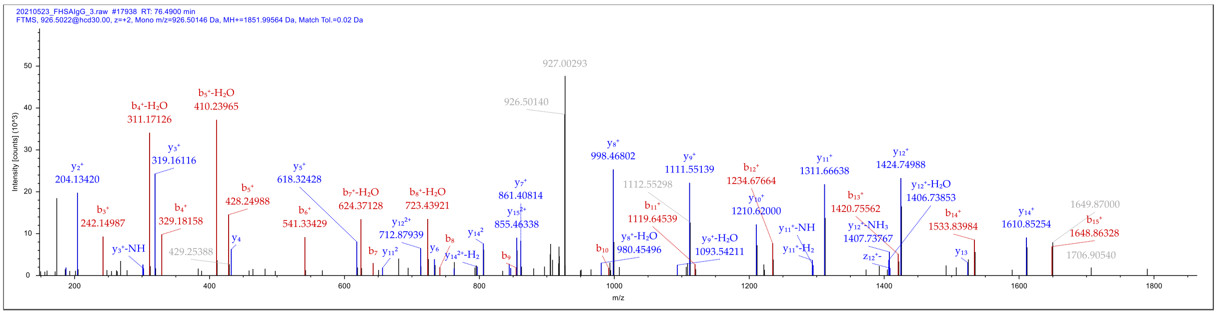


#### AAVSVLTVLHQDWLDGK

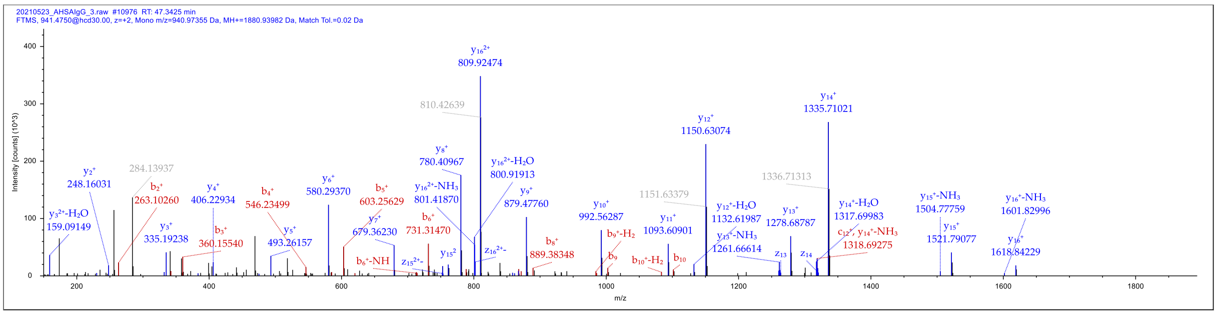


#### FDPWGQGTLVTVSSASTK

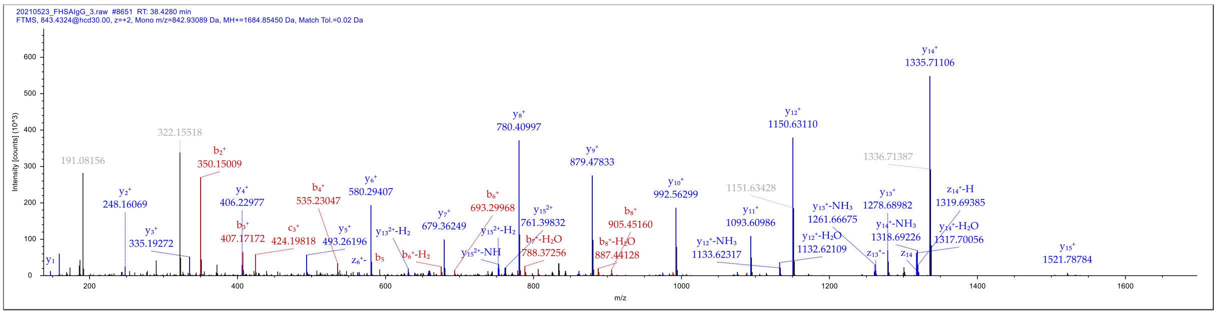


#### YWGQGTLVTVSSASTK

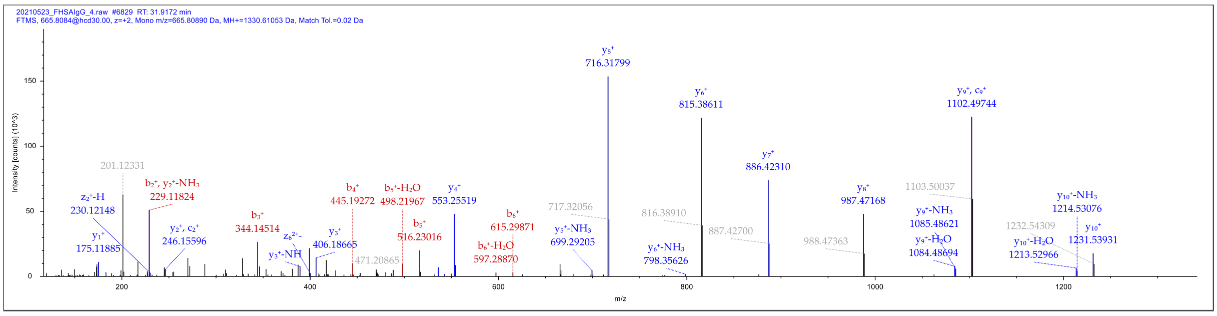


#### VEDTAVYFCAR

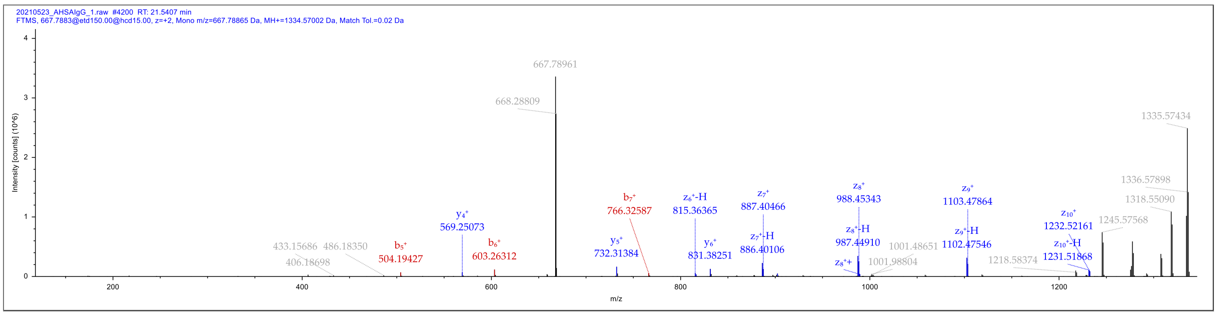


#### SEDTAVYYCAR

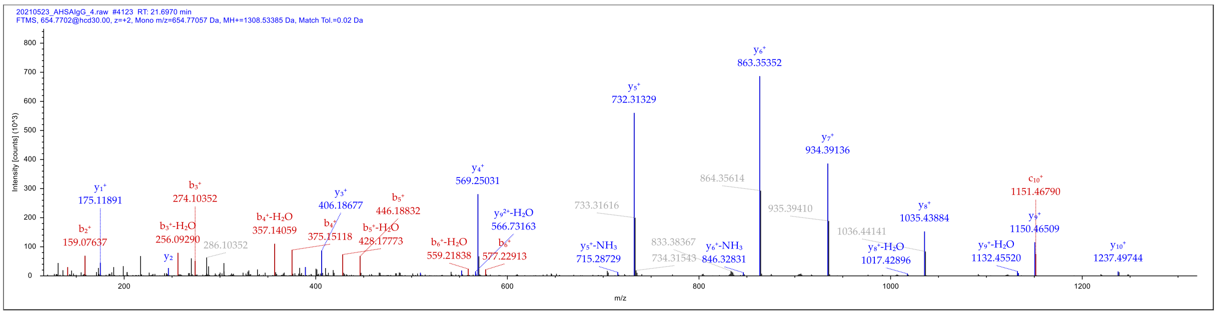


#### ASDTAMYYCAR

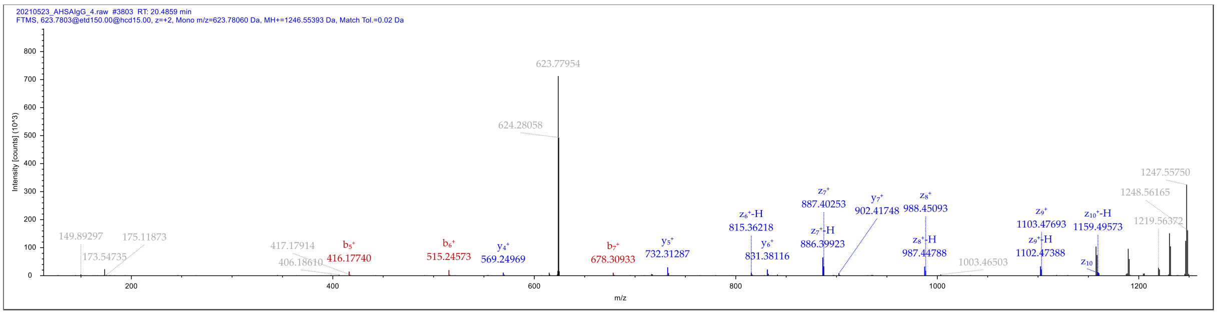


#### AGDTAVYYCAR

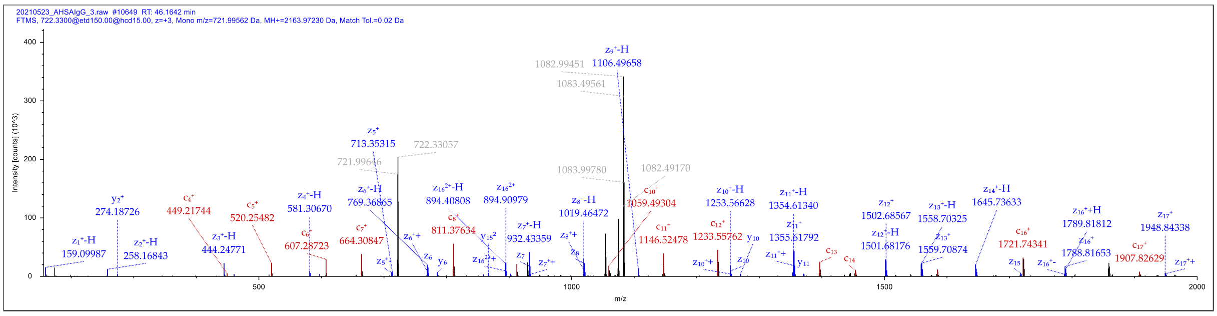


#### LSCAASGFTFSSYGMHWVR

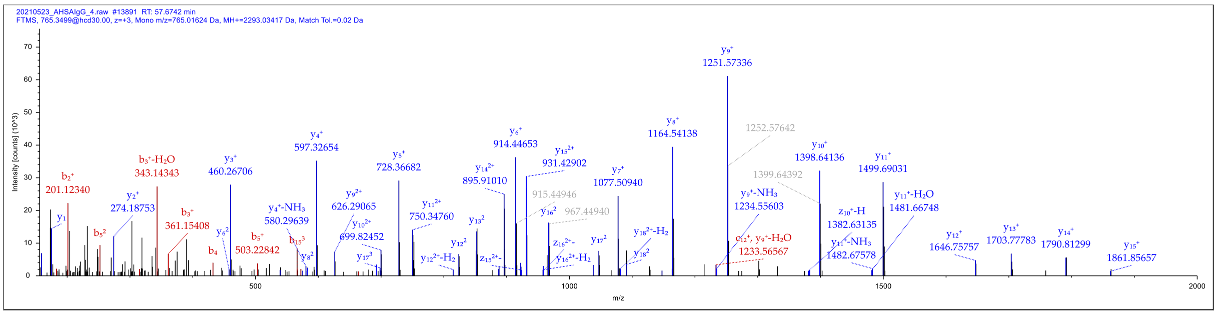

#### LSCAASGFTFSSYWMHWVR

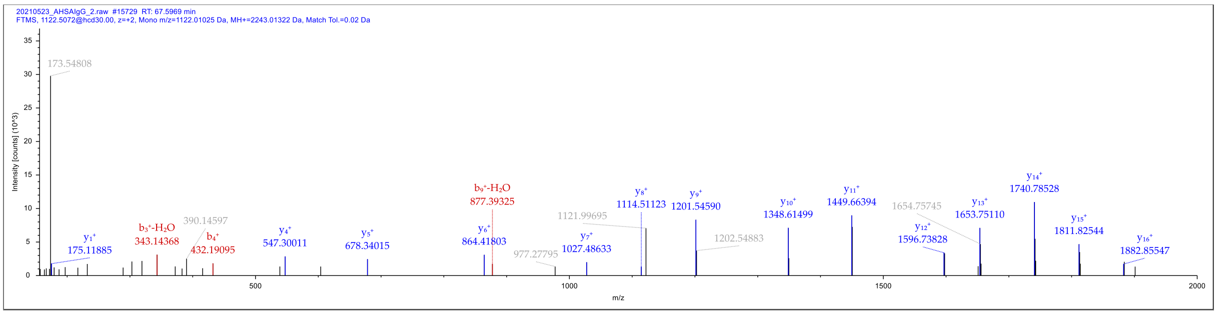


#### LSCAASGFTFSSYWMSWVR

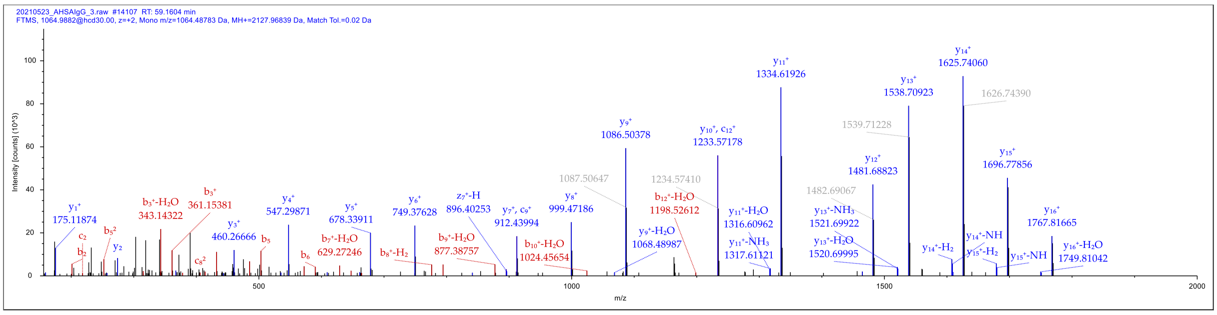


#### LSCAASGFTFSSYAMSWVR

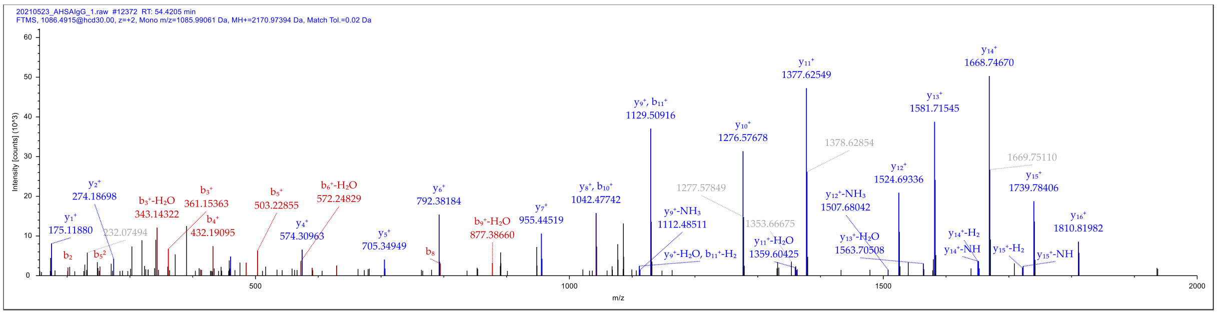


#### LSCAASGFTFSSYSMNWVR

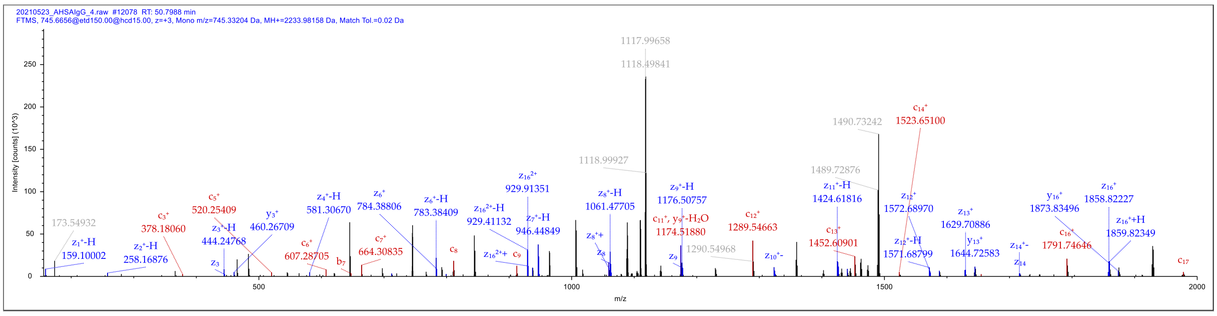


#### LSCAASGFTFDDYAMHWVR

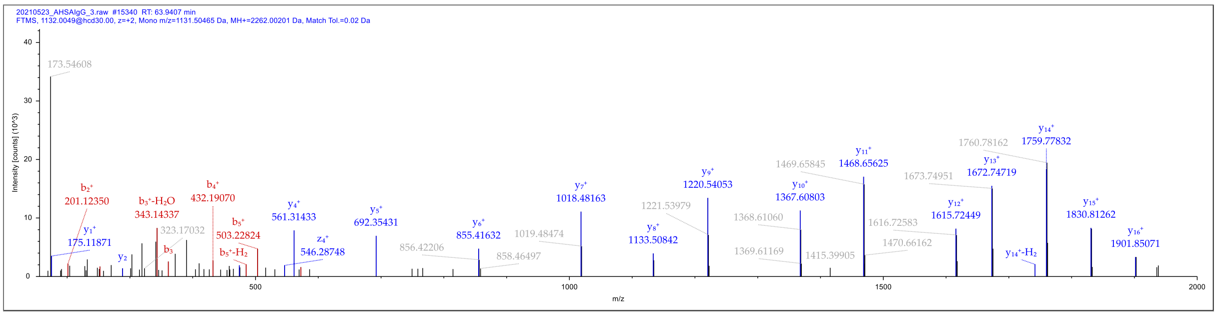


#### LSCAASGFTFSDYYMSWIR

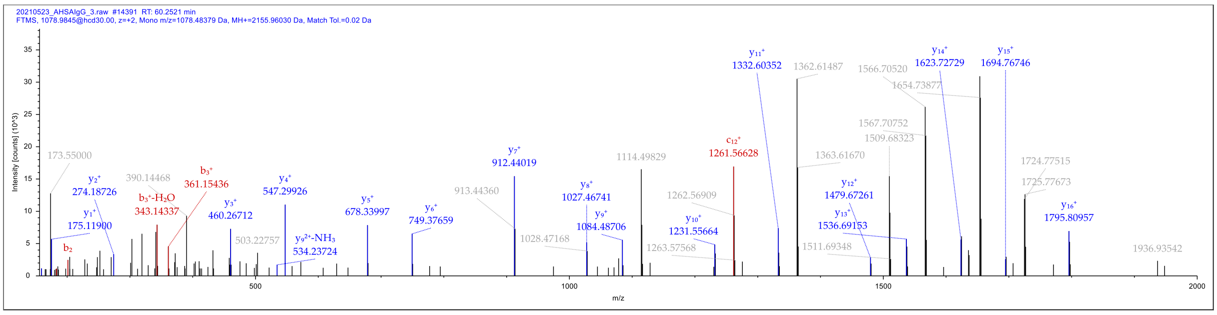


#### LSCTASGFTFGDYAMSWVR

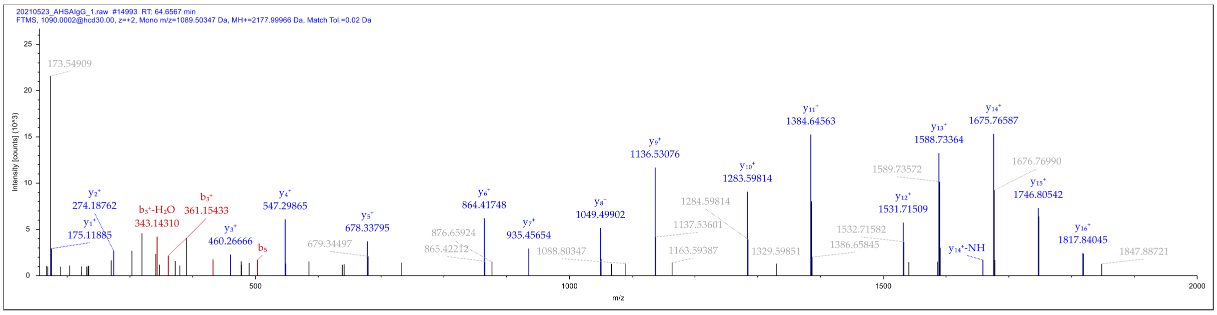


#### LSCAASGFTFSNAWMSWVR

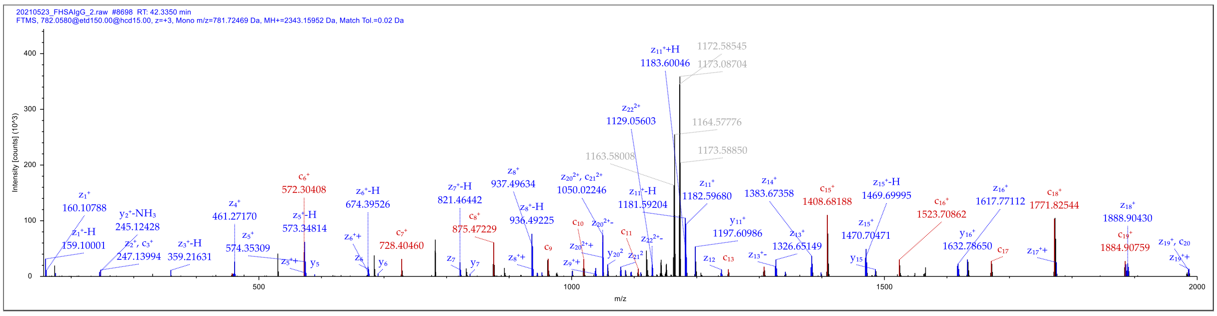


#### ATGIPDRFSGSGSGTDFTLTISR

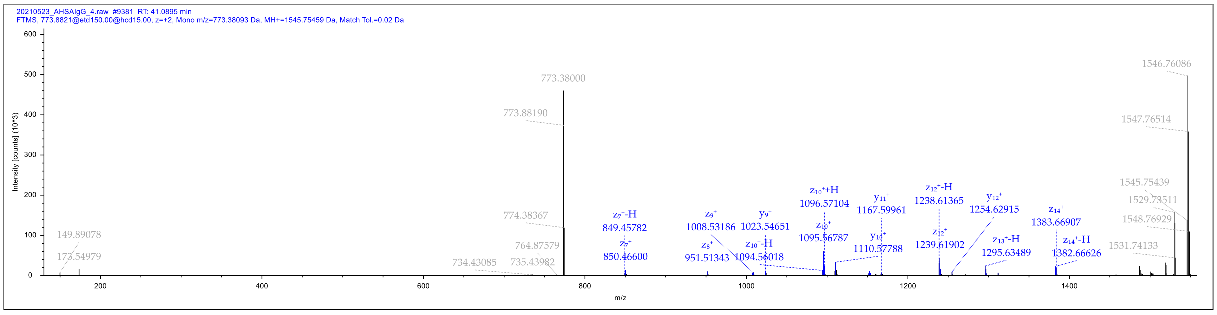

#### FSGSGSGTDFTLTLR

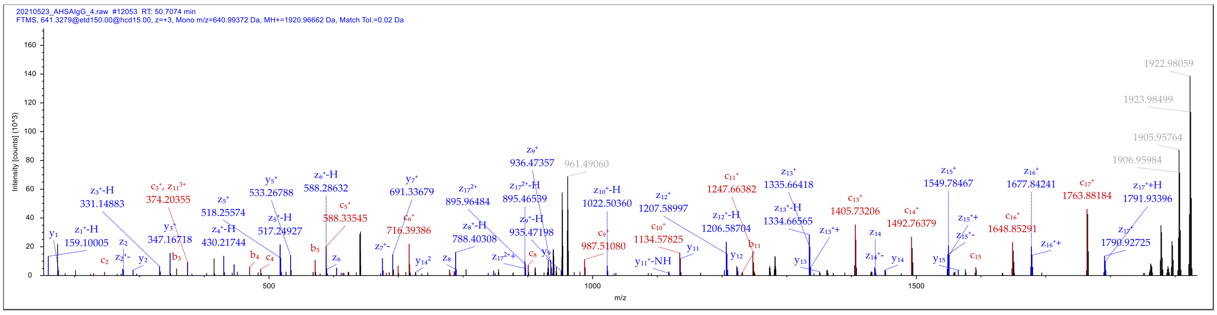


#### DLQLTQSPSFLSASVGDR

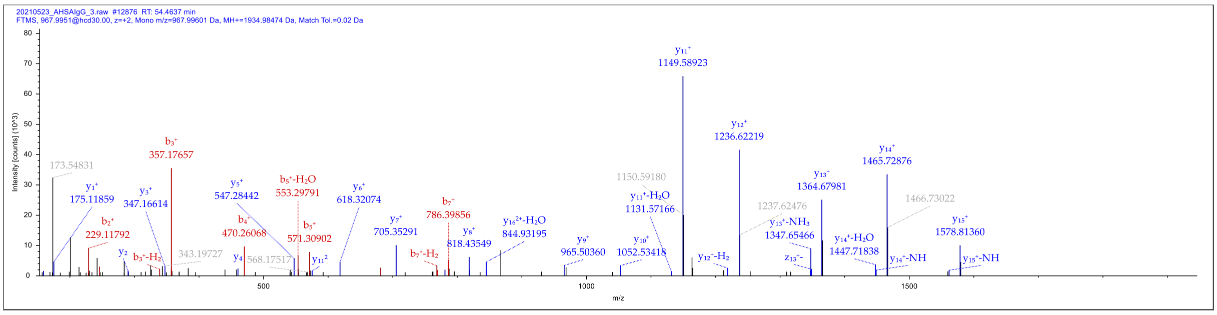


#### DLQLTQSPSFLSASLGDR

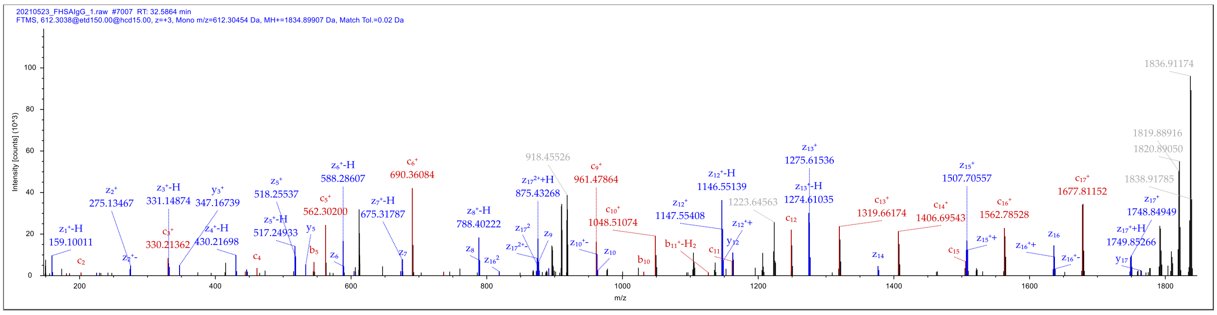


#### ALQMTQSPSSLSASVGDR

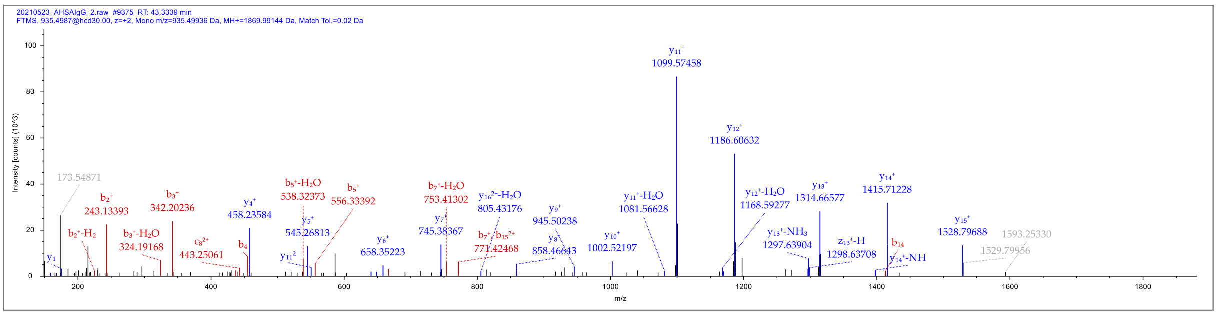


#### ELVLTQSPGSLSLSEGPR

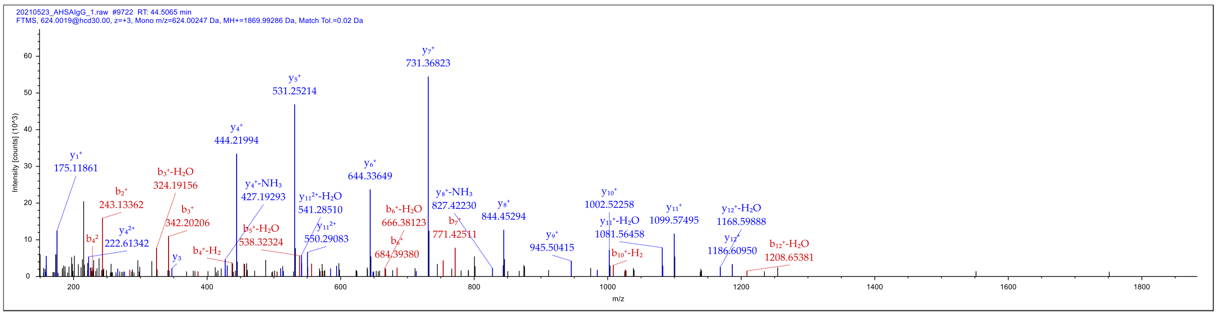


**ELVLTQSPGTLSLSPDGR**

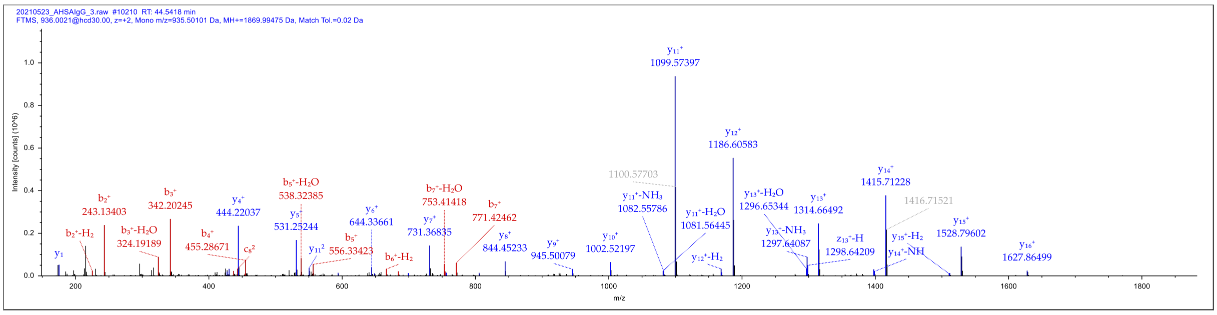


**ELVLTQSPGTLSLSPGDR**

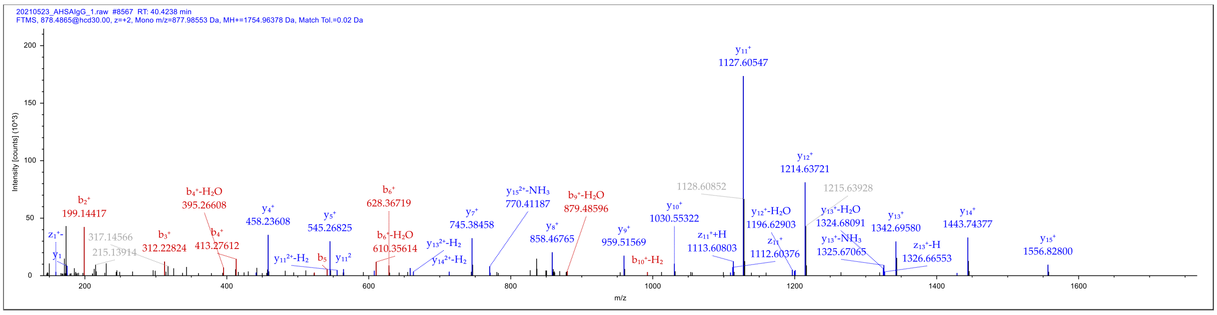


**VVLTQSPATLSLSGPER**

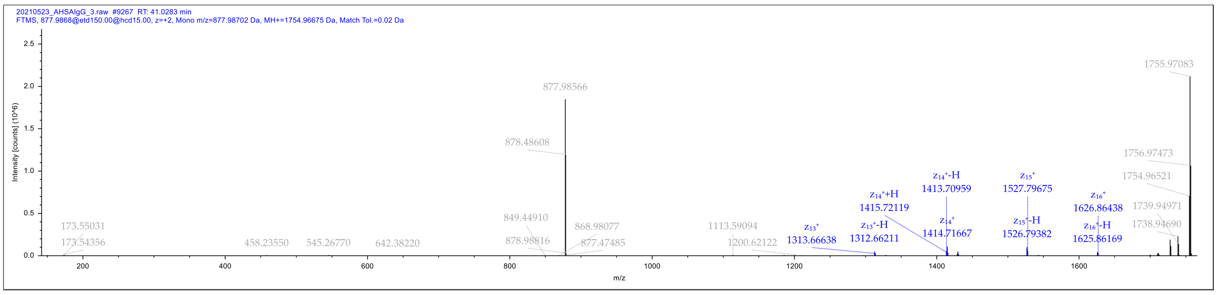


**LVLTQSGPTLSLSPGER**
